# Berberine improves motor deficits in the spastic paraplegia SPG7 mutant mice

**DOI:** 10.64898/2026.06.25.734493

**Authors:** Katarina Paulikova, Anna Sorgente, Emanuela Franchini, Paola Podini, Angelo Quattrini, Linda Pattini, Irene Sambri, Giorgio Casari

## Abstract

Hereditary spastic paraplegia type 7 (SPG7) is a neurodegenerative disorder characterized by progressive motor impairment and cerebellar dysfunction. Mutations in the *SPG7* gene, encoding the mitochondrial metalloprotease paraplegin, disrupt mitochondrial homeostasis and lead to neuronal vulnerability and deficits in motor coordination. Recent studies have identified defective flickering of the mitochondrial permeability transition pore (mPTP) in SPG7 models, suggesting that altered pore dynamics may represent a functional biomarker of mitochondrial dysfunction.

Here, we investigated whether pharmacological modulation of mPTP activity could improve mitochondrial function and motor performance in SPG7 models. Mitochondrial flickering was assessed *in vitro*, while motor behavior was evaluated *in vivo* following chronic treatment with berberine, a natural isoquinoline alkaloid known to modulate mitochondrial bioenergetics. *Spg7*^−/−^ mice and age-matched *Spg7*^+/^ littermate controls received daily oral berberine administration for several weeks, and motor coordination was assessed using the accelerating rotarod test.

Untreated *Spg7*^−/−^ mice exhibited reduced rotarod performance compared with controls, indicating impaired motor coordination. Berberine treatment significantly improved motor performance in pre-symptomatic mutant mice.

These findings indicate that pharmacological modulation of mitochondrial permeability transition pore dynamics can ameliorate motor dysfunction associated with SPG7 deficiency and highlight mPTP flickering as a functional readout of mitochondrial health.

## Introduction

A growing body of evidence indicates that the mitochondrial permeability transition pore (mPTP) plays a central role in mitochondrial physiology (Bernardi et al., 2023) and neuronal survival (Baev et al., 2024). Under physiological conditions, the pore undergoes rapid and transient openings, commonly referred to as mitochondrial flickering, which contributes to the regulation of mitochondrial calcium homeostasis, redox balance, and metabolic signaling (Bernardi et al., 2023; Baev et al., 2024).

These short-lived openings are thought to facilitate controlled ion exchange between the mitochondrial matrix and the cytosol, thereby preventing excessive mitochondrial calcium and ROS accumulation and maintaining cellular homeostasis (Wacquier et al., 2020).

Disruption of this physiological mPTP dynamics has been increasingly linked to neurodegenerative processes (Baev et al., 2024). In hereditary spastic paraplegia type 7 (SPG7), impaired mPTP flickering represents a possible pathogenic mechanism. Loss of paraplegin function disrupts mitochondrial homeostasis, leading to reduced transient pore openings, defective calcium handling, and increased neuronal vulnerability (Sambri et al., 2020; Wali et al., 2023).

These findings establish a mechanistic link between mitochondrial proteostasis defects and altered mitochondrial signaling in SPG7 pathology. Consistent with this model, pharmacological modulation of mPTP function has been shown to rescue mitochondrial defects in SPG7-deficient cellular models (Sambri et al., 2020). More broadly, targeting the pore restores mitochondrial dynamics and improves cellular bioenergetics, supporting mPTP regulation as a promising therapeutic strategy in mitochondrial neurodegenerative disorders.

Among compounds capable of modulating mitochondrial function, the natural isoquinoline alkaloid berberine has attracted increasing interest due to its broad spectrum of biological activities and its ability to regulate mitochondrial metabolism and, more specifically, to increase mitochondrial mPTP flickering (Franchini et al., 2026). Berberine has been widely studied for its metabolic and neuroprotective effects and has been shown to activate AMP-activated protein kinase (AMPK), reduce oxidative stress, and improve mitochondrial bioenergetics in several experimental models (Lu et al., 2015; Cheng et al., 2022). In addition, it promotes mitochondrial quality control pathways, including mitophagy and mitochondrial dynamics, thereby contributing to the maintenance of mitochondrial integrity under pathological conditions (Um et al., 2023).

Given the central role of mitochondrial dysfunction in SPG7 pathogenesis, strategies aimed at restoring mitochondrial homeostasis represent a rational therapeutic approach. In particular, compounds capable of improving mitochondrial signaling and preventing mitochondrial dysfunction may counteract the neuronal vulnerability associated with impaired mPTP dynamics. Within this framework, berberine represents a promising candidate due to its ability to modulate mitochondrial metabolism and enhance cellular resilience to metabolic stress (Franchini et al., 2026;Um et al., 2023).

Based on this rationale, we investigated whether chronic administration of berberine could ameliorate motor deficits in a *Spg7*-null mouse model. By combining *in vitro* measurements of mitochondrial flickering with *in vivo* behavioral assessment using the accelerating rotarod paradigm, we sought to determine whether pharmacological modulation of mitochondrial function can translate into functional improvements in SPG7-associated neurodegeneration.

## Material and Methods

### Cell cultures

Mouse Embryonic Fibroblasts (MEFs) were cultured in Dulbecco’s Modified Eagle Medium (DMEM; Cat. #ECM0728L, EuroClone, Milan, Italy) supplemented with 5% fetal bovine serum (FBS; Cat. #35-079-CV, Corning^®^, Corning Incorporated, NY, USA) and 1% antibiotics (penicillin/streptomycin; PS; Cat. #ECB3001D, EuroClone, Milan, Italy) at 37°C and 5% CO_2_. Cells were split upon reaching confluence by rinsing the monolayer twice with sterile Dulbecco’s phosphate-buffered saline (DPBS; Cat. #21-031-CV, Corning^®^, Corning Incorporated, NY, USA) and incubating with 0.25% Trypsin-EDTA (Cat. #ECM0920D, EuroClone, Milan, Italy) for 3 min. Detached cells were centrifuged to remove trypsin from the medium and then replated.

### Flickering event detection and quantification

mPTP flickering was monitored using a fluorescence-based approach and quantified using a custom MATLAB-based analytical pipeline as previously described (Franchini et al., 2026).

### Mouse model and treatment

All animal procedures were approved by the Institutional Animal Care and Use Committee of the IRCCS San Raffaele Scientific Institute and were conducted in accordance with European Directive 2010/63/EU. S*pg7* mice were generated by *in vitro* fertilization (IVF) using revitalized cryopreserved sperm from the original S*pg7* mouse line (*Spg7*^−/−^) (Ferreirinha et al., 2004) and kept in C57BL/6J strain.

Age-matched *Spg7*^+/−^ mice were used as controls. Mice were housed in groups of up to five animals/cage under 12-hr light/dark cycles, with ad libitum access to food and water. Assignment of animals to treatment groups was conducted in a random manner and sex-balanced. Researchers were blinded to genotype and treatment groups during behavioral test and analysis of data.

*Spg7*^−/−^ knockout (K.O.) and age-matched *Spg7*^+/−^ mice began treatment at either 5 or 8 months of age and received daily oral gavage until they reached 10 or 12 months of age, respectively. Berberine was administered at a dose of 100 mg/kg/day, suspended in a 0.5% sodium carboxymethyl cellulose (CMC-Na) saline solution. Control groups received the same volume of 0.5% CMC-Na saline solution used to dissolve berberine.

### Rotarod performance

Motor performance was evaluated using an accelerating Rota-Rod apparatus (LE8205, PanLab, Harvard apparatus, Barcellona, Spain). All behavioral tests were conducted during the light phase (9:00 AM–6:00 PM) and animals were accustomed to the testing room for at least 30 min on the day preceding testing.

Two experimental cohorts were analyzed: a pre-symptomatic and a post-symptomatic group, which started treatment at 5 and 8 months of age, respectively. Baseline motor performance was assessed in both cohorts prior to treatment. The same animals were subsequently treated daily with berberine (100 mg/kg/day) or vehicle and re-tested at 10 months (pre-symptomatic cohort) or 12 months of age (post-symptomatic cohort).

For each trial, mice were placed on the accelerating rod starting at 5 rpm and reaching 40 rpm over 10 min. Animals underwent five trials per day for three consecutive days, with a maximum trial duration of 300s and at least 15 minutes of rest between each trial. Latency to fall was recorded as the primary outcome measure.

Both male and female mice were included in the analysis. Berberine administration did not induce significant changes in body weight (data not shown). All behavioral assessments were performed by the same experimenter, blinded to genotype and treatment.

### Statistical analysis

Statistical comparisons between two groups were performed using unpaired Student’s t-test or Mann-Whitney U test when normal distribution could not be assumed. Flickering data generated by MATLAB script from each experiment were normalized to the mean of control cell samples or vehicle-treated controls and are reported as scatter dot plots with median lines. Statistical analyses were performed using GraphPad Prism 10. Data were tested for normality and homogeneity of variance using the Shapiro-Wilk normality test. A p-value <0.05 was considered statistically significant. All experiments were performed independently at least three times under identical conditions.

## Results

To investigate mitochondrial membrane potential dynamics in SPG7 deficiency, mitochondrial flickering (mF) events were analyzed in *Spg7* MEFs as already reported (Franchini et al., 2026).

Consistent with previous reports describing impaired mPTP dynamics in SPG7 deficiency [6], *Spg7*^−/−^ MEFs displayed a marked reduction in mitochondrial flickering compared with *Spg7*^+/−^ control cells (Figure 1A).

**Figure 1.**
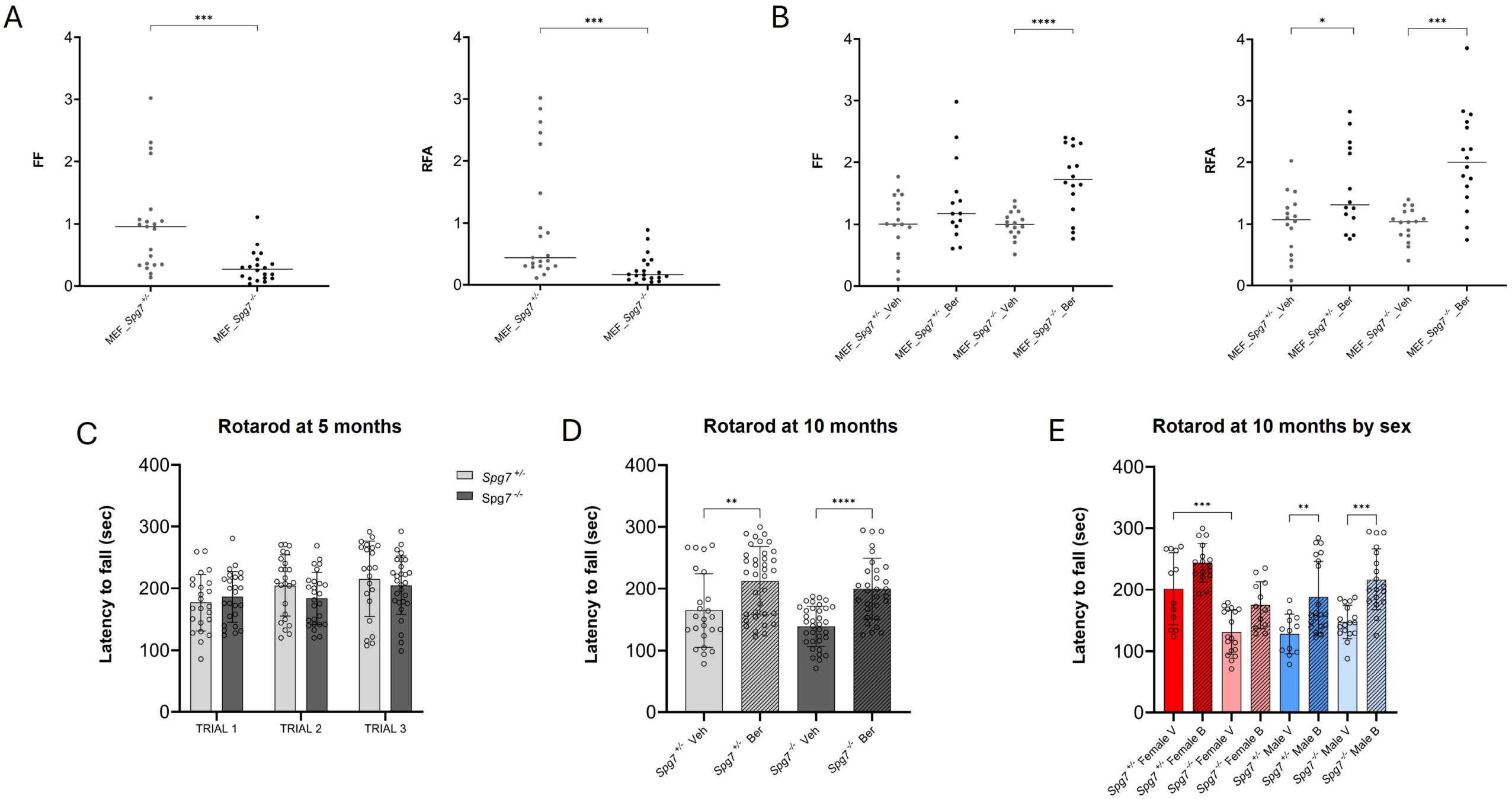
Effects of berberine on mPTP flickering in Spg7 MEFs and on motor performance in Spg7 mice. (A) Basal mPTP flickering in Spg7^+/−^, as control, and *Spg7*^−/−^ MEFs (three independent replicates; Mann–Whitney test, ***p < 0.001). (B) mPTP flickering after berberine treatment (four independent replicates; Welch’s t-test, *p < 0.05, ***p < 0.001, ****p < 0.0001). (C) Motor performance (rotarod test) of untreated Spg7^+/−^ and *Spg7*^−/−^ mice at 5 months of age (Spg7^+/−^ n=26; Spg7^−/−^ n=29). (D) Motor performance of mice pre⍰symptomatically treated with berberine (Ber) or left untreated and assessed at 10 months of age (both sexes combined). (E) As in (D), but with mice analyzed by sex. Ber-treated groups: Spg7^+/−^ females n=6; Spg7^−/−^ females n=5; Spg7^+/−^ males n=7; Spg7^−/−^ males n=7) or vehicle-treated (Veh) (Spg7^+/−^ females n=4; Spg7^−/−^ females n=6; Spg7^+/−^ males n= 5; Spg7^−/−^ males n=6). Dots represent individual mice. Statistical tests (unpaired t-test, Mann–Whitney test, or Welch’s t-test) were applied as appropriate (p < 0.05, **p < 0.01, ****p < 0.0001).

Quantitative analysis normalized to the mitochondrial network area revealed a significant decrease in both flickering frequency (FF; the overall frequency of events per mitochondrial network area) and relative flickering area (RFA; the percentage of mitochondrial area undergoing flickering events per second) in *Spg7*^−/−^ MEFs relative to controls.

Treatment with berberine significantly increased both the frequency of flickering events and relative flickering area, in *Spg7*^−/−^ MEFs compared to vehicle-treated cells, indicating restoration of physiological mPTP activity (Figure 1B).

Next, we investigated whether the restoration of mitochondrial permeability transition pore (mPTP) flickering observed in cellular models could translate into functional improvements *in vivo*. To this end, we assessed the effect of chronic berberine treatment on motor coordination in the mouse model of SPG7 deficiency (Ferreirinha et al., 2004). *Spg7*^−/−^ mice and age-matched *Spg7*^+/−^ littermate controls were randomly assigned to berberine-treated or vehicle-treated groups. Animals received daily oral gavage of berberine (100 mg/kg/day) or vehicle starting at either the pre-symptomatic stage (5 months of age) or the post-symptomatic stage (8 months of age), and treatment continued until the end of the experimental period.

Motor performance was evaluated using the accelerating rotarod test. At baseline, at 5 moths of age *Spg7*^−/−^ mice did not show significant differences in rotarod performance compared with control littermates, confirming absence of SPG7-related symptoms. However, with disease progression untreated *Spg7*^−/−^ mice exhibited a marked reduction in latency to fall, reflecting impaired motor ability (Figure 1C). Chronic administration of berberine significantly improved rotarod performance in the pre-symptomatic cohort, partially rescuing the motor deficit observed in mutant mice (Figure 1D-1E). In the post-symptomatic cohort, berberine-treated animals showed a tendency toward improved rotarod performance relative to vehicle-treated mutants, although the magnitude of the rescue was less pronounced compared with the pre-symptomatic group and not significant (data not shown).

## Discussion

Mitochondrial dysfunction represents a central pathogenic mechanism in hereditary spastic paraplegia type 7 (SPG7), a neurodegenerative disorder caused by mutations in the mitochondrial metalloprotease paraplegin. Previous studies have shown that SPG7 deficiency disrupts mitochondrial homeostasis, leading to impaired mitochondrial dynamics, defective calcium handling, and progressive neuronal degeneration. In particular, impaired mitochondrial flickering has recently emerged as a key player of SPG7 pathomechanism (Sambri et al., 2020).

In the present study, we combined cellular and *in vivo* approaches to investigate whether pharmacological modulation of mitochondrial function could ameliorate disease-associated phenotypes in the SPG7 mouse model. Berberine had previously emerged as one of the most robust modulators of mitochondrial permeability transition pore flickering in a high-content screening of approximately 2,000 FDA/EMA-approved compounds (Franchini et al., 2026), providing a strong rationale for evaluating its therapeutic potential *in vivo*. Using fluorescence-based imaging and a custom analytical pipeline, we confirmed that mitochondrial flickering is significantly reduced in Spg7-deficient MEFs. These findings are consistent with previous studies showing that loss of paraplegin alters mitochondrial permeability transition pore dynamics and compromises mitochondrial signaling pathways involved in calcium homeostasis and metabolic regulation.

Importantly, treatment with berberine significantly increased mitochondrial flickering activity in both control and *Spg7*^−/−^ MEFs, indicating complete restoration of physiological mPTP dynamics. These findings validate previous pharmacological screening results and support the concept that modulation of mitochondrial permeability transition pore activity represents a viable strategy to correct mitochondrial dysfunction associated with SPG7 deficiency.

Beyond cellular phenotypes, we investigated whether pharmacological modulation of mitochondrial function could translate into functional improvements *in vivo*. Behavioral analysis confirmed that *Spg7*^−/−^ mice exhibit progressive motor impairment, consistent with the neurological phenotype previously described in this model. Chronic administration of berberine significantly improved rotarod performance in pre-symptomatic mutant mice, indicating that early intervention can rescue SPG7-induced motor deficits. In contrast, treatment initiated after symptom onset produced only modest improvements, suggesting that the efficacy of mitochondrial-targeted interventions may depend on the timing of treatment.

The beneficial effects of berberine observed in this study may be related to its broader impact on mitochondrial metabolism and cellular energy homeostasis. Berberine has been reported to activate AMPK signaling, improve mitochondrial bioenergetics, and reduce oxidative stress in several experimental systems (Lu et al., 2015). In addition, emerging evidence indicates that berberine can influence mitochondrial quality control pathways, including mitophagy and mitochondrial dynamics. Restoration of physiological mPTP flickering may therefore represent one component of a broader mitochondrial protective effect mediated by this compound.

From a translational perspective, our findings demonstrate that pharmacological restoration of mitochondrial signaling can lead to measurable improvements in motor performance in a model of SPG7 deficiency. The integration of cellular and behavioral analyses strengthens the link between mitochondrial dysfunction and disease phenotype, supporting the concept that mitochondrial permeability transition pore dynamics may serve both as a functional biomarker of mitochondrial health and as a potential therapeutic target in SPG7 and related mitochondrial neurodegenerative disorders.

Some limitations should be considered when interpreting these findings. First, behavioral assessment in this study relied primarily on the rotarod test, which evaluates motor impairment but does not capture the full spectrum of neurological impairments associated with SPG7 pathology. Additional behavioral and neurophysiological analyses may provide a more comprehensive characterization of the therapeutic effects of berberine or possible side effects. Second, while our results demonstrate restoration of mitochondrial flickering in cellular models, further studies will be required to determine whether similar mitochondrial effects occur in affected neuronal populations *in vivo*.

Despite these limitations, the present work provides proof-of-concept evidence that targeting mitochondrial permeability transition pore dynamics represents a promising therapeutic strategy for SPG7-associated neurodegeneration. By linking restoration of mitochondrial flickering to improvements in motor performance, our study highlights the potential of mitochondrial-targeted interventions to modify disease-relevant phenotypes in hereditary spastic paraplegia.

## Acknowledgments

This work was supported by Fondazione Telethon ETS (grant GMR22T2019) and by the Italian Ministry of Health (RF-2019-12370417). We also gratefully acknowledge AB Medica for supporting AS. We acknowledge Alembic, the internal microscopy facility established by IRCCS Ospedale San Raffaele and Università Vita-Salute San Raffaele, Milan, Italy.

## Notes

### Competing Interest Statement

The authors have declared no competing interest.

### Summary of Updates

Two authors have been added. Now the complete list of authors is as follows: Katarina Paulikova1#, Anna Sorgente1#, Emanuela Franchini1, Paola Podini2, Angelo Quattrini2, Linda Pattini3, Irene Sambri4,5 and Giorgio Casari1,2 1 Vita-Salute San Raffaele University, Milan, Italy 2IRCCS Ospedale San Raffaele, Milan, Italy 3Department of Electronics, Information and Bioengineering, Politecnico di Milano, Milan, Italy 4 Telethon Institute of Genetics and Medicine (TIGEM), Naples, Italy 5 CNR-IGM Istituto di Genetica Molecolare Luigi Luca Cavalli-Sforza, Pavia, Italy # These authors contributed equally.

